# Bacterial flagellins: does only size matter?

**DOI:** 10.1101/2020.06.15.151936

**Authors:** Sergei Yu. Shchyogolev, Gennady L. Burygin, Larisa Yu. Matora

## Abstract

The protein sequences of a number of *Azospirillaceae* strains were investigated that contain the pf00669 domain and belong to a variety of bacterial flagellum systems with unusually large flagellin sizes. A set of tools for the homologous and template-based modeling of protein structures was used to reveal the probability of alternative folding, i.e., generation of two protein models with significantly different 3D structures and functions (flagellin or S-layer protein) for a single amino acid sequence.

---

Ref. [1] gives many examples of bacterial flagellar systems with flagellins whose sizes go far beyond those commonly known. It also provides a general overview of their phylogenetic and functional characteristics and estimates their biotechnological potential. In this context, we were interested in the PGPR of the genus *Azospirillum*, which are a popular model for basic and applied studies of associative plant-microbe interactions [2]. When accessing the resource^i^ [1] with an additional filter^ii^, we found (as of May 2020) 232 sequences with sizes up to about 1000 aa, which have the pf00669 domain in their N-terminal portions. When using a set of tools for the homologous and template-based modeling of protein structures for these objects and their homologs, we found proteins that undergo alternative folding. In such cases, two folding variants with considerable differences in protein 3D structure and function (flagellin or S-layer protein) are obtained with high reliability from the same protein sequence. The possibility in principle of the existence of proteins with drastically different 3D structures and with identical amino acid sequences was demonstrated [3] by using the experimental results collected in the Protein Data Bank (PDB)^iii^. Thus, alternative folding can probably be added to the list of various useful and sometimes surprising properties of bacterial flagellins [1]. Then, not only the size but also the specific content of the protein sequence will matter, including the content specifying the degree of intrinsic structural disorder of a protein [4], which will control its conformational flexibility and the possibility of structural transformations in response to various external influences [4, 5].

As an example, we report the data obtained for the hypothetical protein WP_127004468.1 from *Azospirillum doebereinerae* type strain GSF71 with a sequence length of 682 aa, which occupies the seventh position in the list of objects in the resource^ii^. Note that the typical sequence size of the expressed *Azospirillum* full-sized flagellin is 621 aa [6]. As the main tool for predicting 3D protein structures, we used the I-TASSER^iv^ server, supplementing in some cases these results by using the SWISS-MODEL^v^ and MODELLER^vi^ programs. For 3D models of this protein, the I-TASSER program used templates found in PDB^iii^. Of those, the most probable ones were two alternative templates: flagellin from *Salmonella typhimurium* LT2, PDB 1UCU_A (model 1 in Figure 1A) and S-layer protein from *Caulobacter vibrioides* CB15, PDB 5N8P_A (model 2 in Figure 1A). The similarity between the model and its structural analog is characterized by the TM-score^iv^, which had high values for both models in Figure 1A: 0.91 (model 1) and 0.98 (model 2). The TM-score varies within (0-1], where 1 indicates a perfect match between the two structures and the condition TM-score > 0.5 corresponds to the foldings similar in terms of criteria of SCOP/CATH databases. The distribution of regions of the protein intrinsic structural disorder in the sequence WP_127004468.1, as found by several methods [7–9], is shown in Figure 1B (1-3). A comparison of them with the location of the protein secondary structures predicted by I-TASSER (Figure 1B, 4) allows us to suggest the lability of the secondary structures corresponding to these regions and their possible transformations [4, 5] as a probable mechanism of the switching of the folding between the two alternatives described above. Theoretically, alternative folding suggests the presence of two local minima with approximately equal depths in the energy landscape of the protein, which may also depend on the specific content of the protein sequence [10].

**Figure 1.**
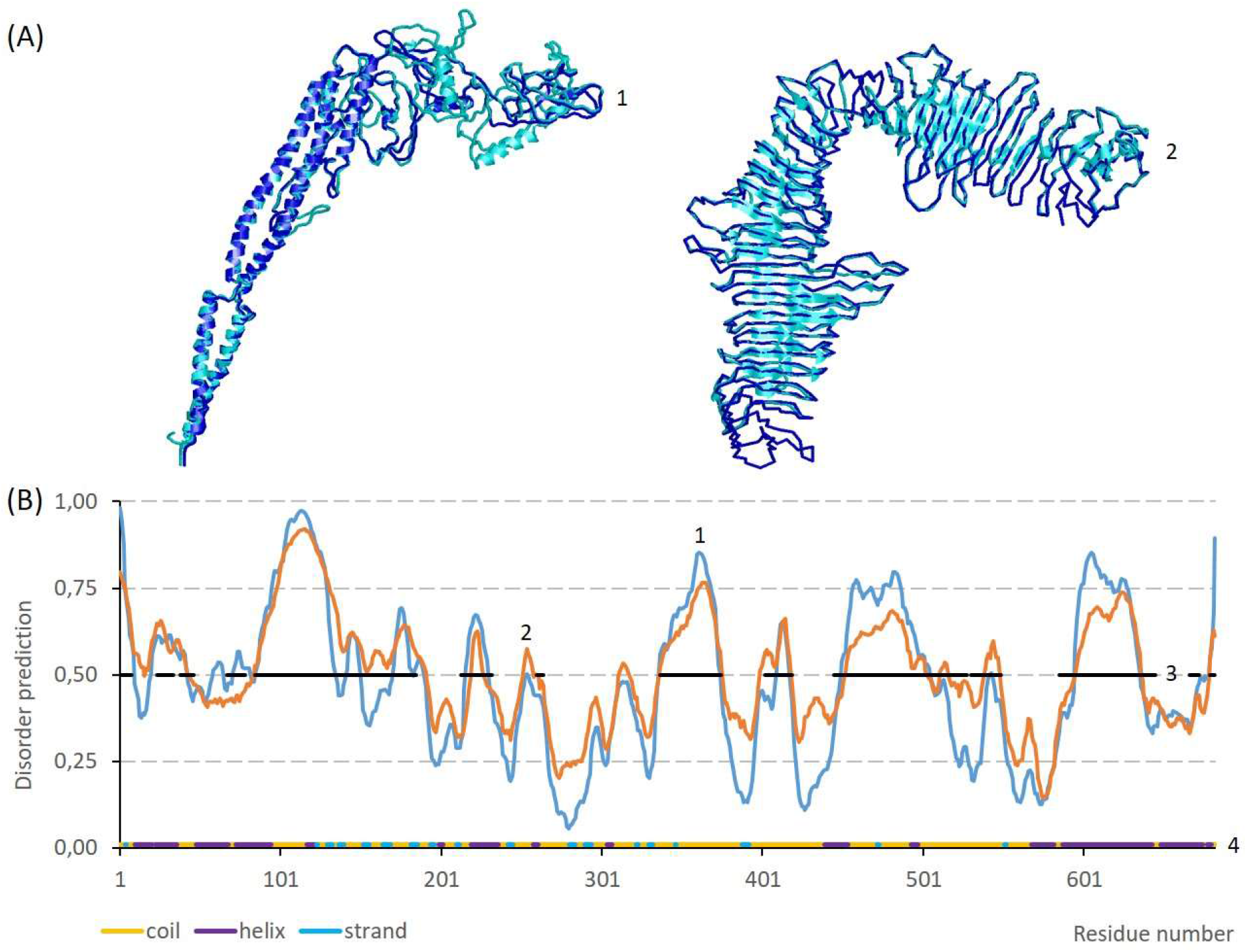
Alternative protein folding for the sequence WP_127004468.1. (**A**) 3D alignment of alternative models (cyan) with their structural analogs from the Protein Data Bank (blue), 1UCU_A (1) and 5N8P_A (2). Visualization with RasMol [11] by using the cartoon (1), (2, cyan) and backbone (2, blue) display options. (**B**) Regions of intrinsic structural disorder in the sequence WP_127004468.1, as predicted with IsUnstruct [7] (1), VSL2 [8] (2) (with disorder prediction > 0,5), and MetaDisorderMD2 [9] (black bar in the middle of the graph) (3), in comparison with the location of the protein secondary structures predicted with I-TASSER (4).

Using Standard Protein BLAST^vii^ with the sequence WP_127004468.1 from *A. doebereinerae* GSF71 versus the non-redundant protein sequences (nr) database, we expanded the set of objects annotated as flagellin-like proteins and showing alternative folding between flagellin and S-layer protein. These include proteins from *A. brasilense* type strain Sp7 (WP_035671652.1), *A. brasilense* Sp245 (WP_014197583.1), *A. lipoferum* 4B (WP_014188288.1), *Nitrospirillum amazonense* CBAmC (WP_088873279.1), *Niveispirillum irakense* DSM 11586 (WP_029012913.1), and some other proteins. The last two strains were previously separated from the genus *Azospirillum* to form independent genera within the family *Azospirillaceae* according to the nomenclature of ref. [12], which develops prokaryote taxonomy on the basis of genome-wide data. Since the publication of ref. [3], the PDB^iii^ has expanded significantly. As a result, the challenge to homologous modeling [3] was overcome in our studies because we used I-TASSER (which is also constantly evolving^viii^) to detect the PDB templates that helped to predict the alternative folding of the flagellin-like proteins that was described above.

In conclusion, we note that the Evolutionary causes and the importance of alternative protein folding, predicted by us for the *Azospirillaceae* flagellins, could be supposedly added to the list of prospects and unresolved questions given in [1].

## Acknowledgments

We thank Dmitry N. Tychinin for his help with the preparation of the English version of this article.

## Resources

i https://www.uniprot.org/uniprot/?query=pf00669&sort=length

ii https://www.uniprot.org/uniprot/?query=pf00669ANDorganism:“azospirillum”&sort=length

iii https://www.rcsb.org

iv https://zhanglab.ccmb.med.umich.edu/I-TASSER

v https://swissmodel.expasy.org

vi https://salilab.org/modeler

vii https://blast.ncbi.nlm.nih.gov/Blast.cgi?PROGRAM=blastp&PAGE_TYPE=BlastSearch&LINK_LOC=blasthome

viii https://zhanglab.ccmb.med.umich.edu/C-I-TASSER

